# When will I get my paper back? A replication study of publication timelines for health professions education research

**DOI:** 10.1101/783332

**Authors:** Lauren A. Maggio, William E. Bynum, Deanna N. Schreiber-Gregory, Steven J. Durning, Anthony R. Artino

**Author notes:** Corresponding author: Lauren A. Maggio, PhD, 4301 Jones Bridge Road Bethesda, MD 20814. Disclaimer: The views expressed in this article are those of the authors and do not necessarily reflect the official policy or position of the Uniformed Services University of the Health Sciences, the Department of Defense, or the U.S. Government.

## Abstract

Biomedical researchers have lamented the lengthy timelines from manuscript submission to publication and highlighted potential detrimental effects on scientific progress and scientists’ careers. In 2015, Himmelstein identified the mean time from manuscript submission to acceptance in biomedicine as approximately 100 days. The length of publication timelines in health professions education (HPE) is currently unknown.

This study replicates Himmelstein’s work with a sample of 14 HPE journals published between 2008-2018. Using PubMed, 19,182 article citations were retrieved. Open metadata for each was downloaded, including the date the article was received by the journal, the date the authors resubmitted revisions, the date the journal accepted the article, and date of entry into PubMed. Journals without publication history metadata were excluded.

Publication history data was available for 55% (n=8) of the journals sampled. The publication histories of 4,735 (25%) articles were analyzed. Mean time from: (1) author submission to journal acceptance was 180.93 days (SD=103.89), (2) author submission to posting on PubMed was 263.55 days (SD=157.61), and (3) journal acceptance to posting on PubMed was 83.15 days (SD=135.72).

This study presents publication metadata for journals that openly provide it—a first step towards understanding publication timelines in HPE. Findings confirm the replicability of the original study, and the limited data suggest that, in comparison to biomedical scientists broadly, medical educators may experience longer wait times for article acceptance and publication. Reasons for these delays are currently unknown and deserve further study; such work would be facilitated by increased public access to journal metadata.

**What this article adds:** Publication delays can negatively impact science and researchers’ careers. A previous study, in biomedicine, leveraged publicly available data to identify a 100-day waiting period between manuscript submission and acceptance. However, this study provides limited information on timelines for health professions education (HPE) articles. Thus, the current study replicates the original investigation with publication timeline data for eight HPE journals, which make their data publicly accessible, finding the mean time from submission to acceptance to be 181 days. Reasons for these delays are currently unknown and deserve further study; such work would be facilitated by increased public access to journal metadata.

---

Researchers have criticized the lengthy timeline from the submission of a manuscript to its ultimate publication, highlighting its detrimental effects to the overall progress of science [1, 2]. This criticism of publication delays may be well-founded. For example, a recent study in *JAMA Oncology* found that results from Phase III oncology trials have a median time to publication of 350 days and even longer if reporting negative findings [3]. While such delays may negatively affect patients, scientists may suffer as well. Researchers have noted that lengthy publication timelines can be detrimental to scientists’ careers, leading to delays of promotion and tenure and/or failure to attain grant funding (e.g., due to scientists’ inability to reference their research under review) [4]. Early career researchers and trainees may be particularly negatively affected [5, 6].

Several studies have investigated publication timelines across a variety of disciplines and publishing models (e.g., open vs. subscription journals, STEM vs. humanities, high vs. low impact factor journals) [7, 8]. A recent *Nature News* investigation and concurrent blog post reported that Himmelstein sampled over 3 million articles from 3,475 biomedical journals present in PubMed with publication metadata between 1965-2015 and found that the average time from submission of a manuscript to its acceptance was approximately 100 days (SD unavailable) [2, 9]. Furthermore, this study found a lag of approximately 25 days between article acceptance and publication in PubMed, which was determined based on data available from 1997-2015. Specific to clinical medicine, another study investigated 781 articles published in 18 internal medicine or primary care journals and identified that the average time from submission to acceptance was 153 days (median=123) with an average lag between acceptance and publication of 105 days (median=68) [10]. While these studies provide two valuable benchmarks for understanding publication timelines across biomedicine and within a clinical discipline, we currently know very little about publication timelines in the field of health professions education (HPE).

Why does assessing for publication delays matter?

We believe that lengthy publication timelines, if present, may be potentially problematic in HPE research; this underlies the impetus for the current study. Indeed, the cognitive, physical, and psychological challenges of learning medicine in demanding clinical environments should be met with timely, up-to-date, and evidence-based educational knowledge and instructional strategies. Lengthy publication timelines may undermine the effectiveness of teachers who strive to deliver evidence-based content in an evidence-based manner within an optimal learning environment. Failure to disseminate the evidence that drives each of these goals may ultimately negatively impact medical learners and, indirectly, patients who rely on learners for competent care. Moreover, publication delays for work that explores important phenomena such as learner suicide or depression may negatively affect medical learners through delays in the implementation and evaluation of novel resources and support structures. In addition, HPE scientists whose work is embargoed during the publication process are unable to receive credit for pending publications in grant applications and subsequent research studies. Finally, long embargoes can hurt researchers whose work in fast-moving areas like educational technology and social media may no longer be contemporary or relevant by the time it is published.

In light of these concerns, we aimed to address the lack of data about publication timelines by replicating Himmelstein’s previous work to examine data in the HPE journals that openly provide this information. While Himmelstein’s work included articles published broadly in biomedicine from 1965-2015, we focus on HPE articles published 2008-2018. We believe this timeframe is appropriate to understand publication timelines in the rapidly growing field of HPE, as only two HPE journals provided publication history data prior to 2010. As such, Himmelstein’s prior work provides limited inclusion of HPE information. We believe these data could be a first step towards addressing potential publication delays in HPE and may spark conversations about publication timelines and ways to optimize them.

## Method

To calculate publication timelines in HPE, we replicated the bibliometric approach reported in *Nature News* by Himmelstein [2, 9]. We chose to replicate this particular approach because, unlike other studies reported in the literature [7, 8, 10], which relied on humans to extract the relevant data, Himmelstein’s approach utilized computer code and publicly available data. Using a computer-based rather than human-powered approach allowed us to more efficiently and objectively extract a large volume of data from articles published between 2008-2018 and to mitigate the risk of human coding errors.

We conducted this replication study with a sample of 14 journals that have been previously identified as core HPE journals [11,12]. Journals included: *Academic Medicine, Advances in Health Sciences Education, BMC Medical Education, Canadian Medical Education Journal, Clinical Teacher, International Journal of Medical Education, Journal of Advances in Medical Education and Practice, Journal of Graduate Medical Education, Medical Education, Medical Education Online, Medical Teacher, Perspectives on Medical Education, Teaching and Learning in Medicine*, and *The Journal of Continuing Education in the Health Professions*. These journals were included in Himmelstein’s original study if they provided publication history metadata between 1965-2015. Thus, this study extends and replicates his earlier work by adding three additional years of data, during which the number of HPE publications steadily increased.

Following Himmelstein’s steps [2,9], we queried PubMed on April 10, 2019 for articles published in these 14 journals between 2008-2018. Our search yielded 19,182 citations, and we downloaded the complete, publicly accessible metadata for each citation. From this metadata, which was generated by National Library of Medicine staff, we identified the journals that make publicly available their article-level publication history (e.g., the timelines for each of the steps in the publication process). This history includes the date the article was received by the journal, the date the authors resubmitted revisions, the date the article was accepted by the journal, and the date that the article was entered into PubMed. Based on this available data, we defined the following three time periods: 1) Publication Time: the time from article submission to appearance in PubMed, 2) Acceptance Time: the time from article submission to acceptance by the journal, and 3) Processing Time: the time from article acceptance to appearance in PubMed. These three time periods aligned with those defined in Himmelstein’s work, and, in similar fashion, we excluded from our analysis journals that did not supply this publication history metadata.

We used SAS 9.4 for analysis and data management. We ran two 2-sided, independent sample t-tests to determine any potential differences in publication timelines based on funding source. To increase the transparency of our work and encourage further replication, we have deposited our data set and corresponding computer code here: https://github.com/DNSchreiber-Gregory/Publication-Timelines/tree/DNSchreiber-Gregory-Publication-Timelines

## Results

Over the course of the study period (2008-2018), 19,182 articles were published in the 14 HPE journals sampled. Of these journals, publication timeline metadata was available for eight of the journals (*Advances in Health Sciences Education*, *BMC Medical Education*, *International Journal of Medical Education, Journal of Advances in Medical Education and Practice, Journal of Graduate Medical Education*, *Medical Education, Medical Education Online,* and *Perspectives on Medical Education)*. During the study period, these eight journals published 8,681 articles. Of these articles, publication history data was available and extracted from 4,735 (55%) articles.

The mean publication time from author submission to posting on PubMed was 263.55 days (SD=157.61; median=228). The mean acceptance time from author submission to journal acceptance was 180.93 days (SD=103.89; median=163). The mean processing time from acceptance by the journal to posting on PubMed was 83.15 days (SD=135.72; median=23). Table 1 presents publication, acceptance and processing times for articles published between 2008 and 2018 in these eight HPE journals.

**Table 1:** Publication, acceptance and processing times, as expressed in days, between 2008 and 2018, in eight HPE journals with available publication timeline metadata.

| Publication Date | Total Articles | Articles w/ timeline metadata (%) | Publication Time |  | Acceptance Time |  | Processing Time |  |
| --- | --- | --- | --- | --- | --- | --- | --- | --- |
|  |  |  | Median | Mean (SD) | Median | Mean (SD) | Median | Mean (SD) |
| **2008** | 993 | 119 | 185 | 203.83<br>(120.20) | 156 | 169.19<br>(88.56) | 7 | 34.64<br>(87.16) |
| **2009** | 1328 | 152 | 172 | 155.82<br>(80.82) | 153.5 | 172.05<br>(77.98) | 7 | 16.23<br>(22.78) |
| **2010** | 1578 | 275 | 199 | 267.16<br>(181.55) | 138.5 | 149.43<br>(84.20) | 29 | 118.07<br>(187.61) |
| **2011** | 1449 | 273 | 229 | 349.41<br>(257.16) | 157 | 158.71<br>(87.47) | 21 | 190.70<br>(246.11) |
| **2012** | 1567 | 286 | 204 | 315.75<br>(253.26) | 162 | 163.91<br>(84.78) | 19 | 155.37<br>(241.28) |
| **2013** | 1796 | 467 | 233 | 275.44<br>(180.55) | 155 | 162.12<br>(96.97) | 25 | 113.32<br>(154.02) |
| **2014** | 1943 | 643 | 260 | 286.67<br>(155.48) | 157 | 172.92<br>(97.60) | 35 | 114.00<br>(138.40) |
| **2015** | 1927 | 609 | 236 | 244.38<br>(120.98) | 157 | 177.45<br>(106.49) | 30 | 67.35<br>(76.88) |
| **2016** | 2126 | 651 | 199 | 227.14<br>(117.55) | 154 | 176.94<br>(106.70) | 18 | 50.19<br>(75.36) |
| **2017** | 2153 | 610 | 250.5 | 256.09<br>(98.98) | 186.5 | 205.58<br>(101.11) | 20 | 50.51<br>(57.44) |
| **2018** | 2322 | 647 | 251 | 265.38<br>(118.32) | 199 | 223.53<br>(124.57) | 20 | 42.21<br>(44.43) |
| **Overall Study Period** | 19182 | 4735 | 228 | 263.55<br>(157.61) | 163 | 180.93<br>(103.89) | 23 | 83.15<br>(135.72) |

### Journals

Reporting of publication history data varied by journal. For example, *BMC Medical Education* and the *International Journal of Medical Education* reported publication history data for 99% of articles published in the study period, whereas *Perspectives on Medical Education* provided publication timeline data for 25% of articles (see Table 2). As noted above, the journals in our analysis made article publication history metadata available to varying degrees, with only four of the journals (*Advances in Health Sciences Education*, *BMC Medical Education, International Journal of Medical Education, and Medical Education Online)* making the data available for more than 50% of their articles (see Table 2). Additionally, in some cases, metadata was only available for certain years of the observed time period (see Figure 1). For example, data were available for *Medical Education Online* between 2010-2016. Only *Advances in Health Sciences Education* and *BMC Medical Education* featured timeline metadata for the entire study period. Of note, while *Medical Education* reported timeline metadata for eight years, the journal reported publication, acceptance and processing times as zero days for two of those years.

**Table 2:** Publication, acceptance, and processing time, expressed in days, by journal for articles between 2008 and 2018.

| Journal Name | Total Articles published | Articles w/ timeline metadata (%) | Publication Time |  | Acceptance Time |  | Processing Time |  |
| --- | --- | --- | --- | --- | --- | --- | --- | --- |
|  |  |  | Median | Mean (SD) | Median | Mean (SD) | Median | Mean (SD) |
| *Advances in Health Sciences Education* | 809 | 769 (95) | 182 | 190.34 (103.76) | 157 | 164.04 (98.17) | 18 | 26.30 (42.24) |
| *BMC Medical Education* | 2077 | 2054 (99) | 217 | 238.51 (114.20) | 203 | 223.56 (112.23) | 11 | 14.91 (14.84) |
| *International Journal of Medical Education* | 299 | 298 (99) | 177 | 196.06 (106.97) | 146 | 161.41 (92.10) | 21 | 35.88 (54.33) |
| *Journal of Advances in Medical Education & Practice* | 162 | 57 (35) | 374.5 | 377.71 (141.83) | 123.5 | 138.65 (75.13) | 241 | 246.72 (147.92) |
| *Journal of Graduate Medical Education* | 1621 | 635 (39) | 506 | 495.58 (197.12) | 158 | 161.78 (67.84) | 337 | 337.29 (187.26) |
| ***Medical Education*** | 2766 | 607 (22) | 275 | 286.97 (87.32) | 130 | 137.85 (64.77) | 141 | 148.79 (66.57) |
| ***Medical Education Online*** | 491 | 305 (62) | 126 | 133.63 (67.21) | 80 | 88.19 (62.50) | 39 | 45.17 (27.83) |
| ***Perspectives on Medical Education*** | 456 | 10 (2) | 206 | 210.70 (119.73) | 159.5 | 163.80 (122.84) | 50 | 46.90 (15.52) |

Publication timeline data was unavailable for *Academic Medicine* (n=4852), *Canadian Medical Education Journal* (n=230), *Medical Teacher* (n=3303), *Teaching and Learning in Medicine* (n=597), *The Journal of Continuing Education in the Health Professions* (n=515), and *The Clinical Teacher* (n=1004).
Of note, none of the included journals provided complete publication metadata, which must be taken into consideration when examining timelines for individual journals.

**Figure 1:**
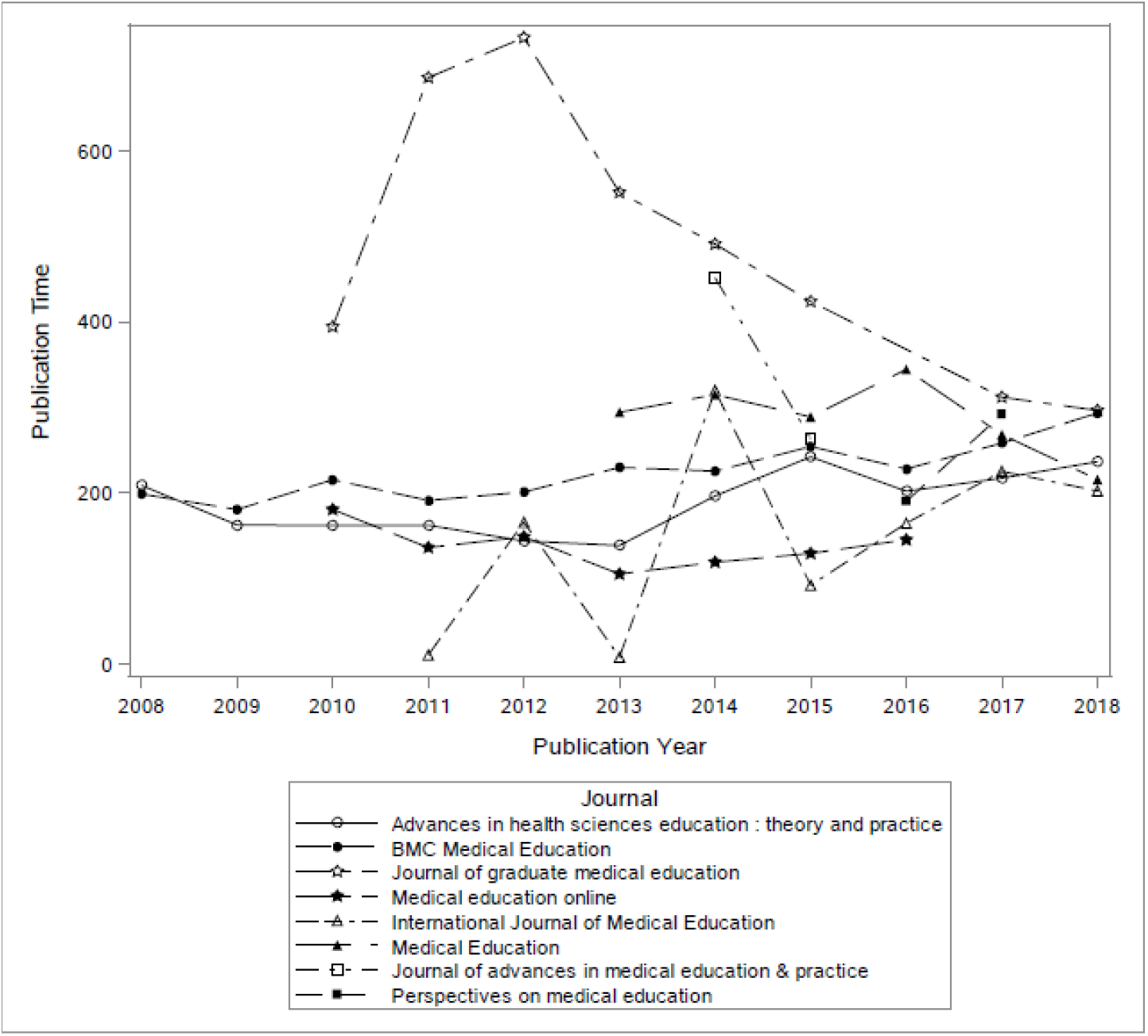
Publication time (i.e. the time from article submission to appearance in PubMed) by journal expressed in days for articles published in HPE journals between 2008 and 2018.

### Publication Types

Articles represented a variety of publication types as indexed by the National Library of Medicine (Table 3). Editorials, which do not typically include revisions, had the shortest publication time of 83.53 days (n=43; SD=97.41; median=33) in contrast with meta-analyses, which had the longest publication time of 279.93 days (n=30; SD=115.88; median=274).

**Table 3:** Publication, acceptance, and processing times, as expressed in days, by publication types featuring 10 or more articles

| Publication Type* | Total Articles | Publication Time |  | Acceptance Time |  | Processing Time |  |
| --- | --- | --- | --- | --- | --- | --- | --- |
|  |  | Median | Mean (SD) | Median | Mean (SD) | Median | Mean (SD) |
| **Clinical Trial** | 15 | 232 | 274.13 (161.04) | 227 | 261.53 (158.80) | 11 | 12.60 (8.13) |
| **Comparative Study** | 259 | 223 | 241.66 (111.54) | 196 | 213.75 (109.73) | 16 | 27.91 (39.07) |
| **Editorial** | 43 | 33 | 83.53 (97.41) | 3 | 47.24 (89.77) | 22.5 | 32.24 (47.07) |
| **Evaluation Studies** | 102 | 211 | 230.84 (125.23) | 167.5 | 199.51 (127.84) | 12 | 31.33 (49.71) |
| **Historical Article** | 20 | 177 | 194.10 (134.71) | 134.5 | 140.00 (125.99) | 26.5 | 54.10 (67.42) |
| **Letter** | 36 | 104 | 147.39 (124.92) | 80.5 | 112.33 (110.15) | 22 | 35.06 (48.73) |
| **Meta-Analysis** | 30 | 274 | 279.93 (115.88) | 184 | 208.27 (127.78) | 28 | 71.67 (68.95) |
| **Multicenter Study** | 82 | 261.5 | 261.55 (115.94) | 198.5 | 206.91 (107.91) | 21 | 54.63 (73.11) |
| **Observational Study** | 48 | 230 | 250.27 (131.16) | 157 | 194.44 (125.21) | 25.5 | 55.83 (65.97) |
| **Randomized Controlled Trial** | 227 | 237 | 241.39 (111.67) | 170 | 196.43 (111.13) | 19 | 44.96 (58.25) |
| **Review** | 258 | 232.5 | 244.49 (103.35) | 161.5 | 175.21 (86.26) | 28.5 | 69.28 (76.69) |
| **Validation Studies** | 109 | 225 | 243.81 (112.28) | 189 | 201.73 (88.33) | 17 | 42.07 (83.93) |
| **Journal Article** | 4646 | 230 | 266.22 (156.88) | 165 | 182.82 (102.96) | 23 | 83.94 (136.53) |

\*Articles can be indexed with more than one publication type.

### Funding

Twenty-one percent of articles (n=1,011) with available publication timeline data reported receiving funding: 11.2% of articles reported receiving funds from the United States (US) government, of which 8.7% (n=88) received funds from the National Institutes of Health; the remainder (92%) reported funding from non-US government sources. When considering these percentages, it is important to note that articles can and often do report multiple funders. When comparing funded versus unfunded research, we observed significant differences in processing time (p < .0001, Cohen’s d = 0.46) and publication time (p < .0001, Cohen’s d= 0.34), with unfunded projects having significantly longer timeframes in both (see Table 3). We did not find a significant difference in acceptance time (p = 0.3074) between funded and unfunded projects.

**Table 3:** Publication, acceptance and processing times, as expressed in days, by reported funding.

| Funding | Total Articles | Publication Time |  | Acceptance Time |  | Processing Time |  |
| --- | --- | --- | --- | --- | --- | --- | --- |
|  |  | Median | Mean (SD) | Median | Mean (SD) | Median | Mean (SD) |
| No Funding Reported | 3720 | 236 | 273.81 (167.82) | 163 | 180.16 (105.43) | 24 | 94.29 (148.01) |
| Total Funded Articles | 1011 | 209 | 225.83 (104.08) | 163 | 183.77 (98.06) | 16 | 42.15 (58.71) |
| Non-US Government Funding | 932 | 211 | 228.02 (104.37) | 166 | 186.40 (98.12) | 16 | 41.72 (58.43) |
| US Government Funding | 114 | 183.5 | 134.44 (114.34) | 132.5 | 173.33 (104.36) | 22 | 38.89 (54.14) |
| National Institutes of Health Funding | 88 | 183 | 195.56 (98.88) | 129.5 | 160.33 (102.16) | 20 | 35.23 (46.82) |
| US Gov't Funding (Non-NIH) | 37 | 200 | 224.51 (100.76) | 154 | 174.27 (89.70) | 29 |  |

\*Articles can report multiple funders.

## Discussion

We have replicated and applied a previous study design within the field of HPE, and our findings suggest that, when compared to the previous study [9], which was broadly focused on 3,475 biomedical science journals with publication history metadata in PubMed, HPE may have longer publication timelines. To our knowledge, these findings represent the first and only available indicators of publication timelines in HPE. However, before discussing these findings and their implications, it is important to address the limitations of our approach.

Similar to the original study [9], our replication analysis was constrained by the incomplete publication history data made publicly available by the journals and their publishers. In our analysis, we were able to analyze only 25% of HPE articles published between 2008-2018. Furthermore, only eight of the 14 journals in our sample made data available, and none provided complete data for all of their articles. The incompleteness of the data may have skewed the results of our analysis, and this should be taken into consideration when examining the timelines for individual journals. Accordingly, we had to exclude some HPE journals, including *Academic Medicine*, which annually publishes the greatest number of articles in the field. Furthermore, our findings represent only a coarse quantitative indicator of publication, acceptance, and processing time. In other words, we were unable to explore the publication process in a fine-grained manner, a process that includes multiple steps and multiple stakeholders (e.g. authors, reviewers, editors, and publishers), each of whom plays a role in decreasing (or extending) publishing timelines. Thus, the data are incomplete, and we are unable to draw detailed conclusions about what exactly occurs within our observed timeframes. However, despite these limitations, we believe our findings draw attention to the potential presence of publication delays and present an opportunity to spark conversation among authors, editors, reviewers and publishers in the HPE community.

Furthermore, our findings support the need for all HPE journals to publish timeline data. Indeed, if there is power in data, there is even greater power in *open-access* data and *open-sourced* data analysis [13, 14]. Unfortunately, despite our ability to conduct this study using publicly available data and a previously developed analytic method, our analysis was hampered by the lack of a complete dataset from all queried journals. Moving forward, we believe HPE journals should compile and make available the data necessary to thoroughly understand the processes governing the publication of our science and to which we are beholden. In order to develop a complete dataset, we call on all journals in HPE to make their publication timeline data publicly available in its entirety. Doing so will promote transparency and help identify the ways in which our publication timelines might be improved.

Awareness of accurate publication timelines in HPE education could benefit numerous stakeholders. For journal editors and publishers, analysis based on complete data would provide an opportunity for benchmarks within the field, critical reflection on their own timelines, and sharing of best practices from exemplars in the field. Journals that use this data to streamline their publication timelines may experience higher submission rates for higher quality articles from authors seeking a faster, more transparent publication process. At the program level, graduate programs in HPE increasingly require students to have publications accepted by peer-reviewed journals in order to graduate [15]. Knowledge of journal timelines may assist in the planning of educational programs and forecasting of graduation timelines. For clinical trainees seeking to publish scholarship to assist residency or fellowship applications, knowing which classes of articles are published more quickly may lead to more timely citations and enhanced applications. For authors, programs, reviewers and editors alike, more thorough data analysis is needed to determine whether variables such as funding sources influence publication timelines and unlevel the playing field for researchers whose work is unfunded.

We believe that such transparency within the publication process is very important for authors. Currently, authors who submit manuscripts to HPE journals lack awareness of the timelines that govern the publication process. We believe this lack of information undermines authors’ ability to be critical about when and where to submit their manuscripts, a decision that may have real implications, especially for those authors facing funding or promotion deadlines or striving to publish contemporary research. Even more fundamental, we believe that transparency about publication timelines should be a basic courtesy afforded to all authors.

While our data are incomplete and do not include data from several key journals in our field, they do suggest that publication timelines – from the journals for which we have meaningful data – are over twice as long as the 100-day benchmark published by Himmelstein [2, 9] and longer than the timelines observed in internal medicine and primary care journals [10]. Thus, lengthy publication delays may be a challenge that needs to be confronted in the HPE community.

### Potential Solutions

While we await greater clarity in the scope and nature of this challenge, which will be aided by consistent data sharing from all our journals, we have identified potential, immediate solutions—at the researcher, editor, and publisher levels—that might be explored to improve existing publication timelines.

#### Researchers

While researchers are bound to the systems that govern publication of their research, they are not powerless to affect meaningful change in publication timelines. Aligning submissions with journals likely to publish them, submitting revisions in a timely manner, strategically avoiding suboptimal submission dates (e.g. prior to major holidays), and following up with journal editorial staff in the event of a delay are active measures researchers can take to facilitate expeditious publication timelines. Researchers might consider disseminating their work via alternate mechanisms while awaiting journal review; such outlets include preprint servers (e.g. bioRxiv or medRxiv) and presenting at professional meetings. That said, authors should be aware of ethical rules related to dual publication and should always disclose such dissemination efforts to editors in their cover letters. Further, when serving as peer reviewers, researchers should make every effort to complete their reviews on time.

#### Journals and editorial staff

Journals and their editorial staff can also act to streamline publication timelines. By allowing – or even encouraging – submission of preprints [16], journals and publishers will facilitate dissemination of science during the peer-review process. For example, articles that appear first as preprints in the life sciences have recently been shown to have a 1.31 increase in citations and higher altmetric attention scores once the articles are published in peer-reviewed outlets [17]. Editors might also consider alternate peer-review approaches likely to streamline time to publication. For example, the post-publication peer-review process that *MedEdPublish* utilizes allows authors to submit an article prior to peer-review, thus providing authors a platform for immediately sharing their research while waiting for invited reviewers to post comments. Alternately, journals such as *Advances in Health Sciences Education* now utilize a “Fast Track” option that allows authors to submit peer reviews from other journals that have previously rejected the submission under review [18].

#### Publishers

Finally, publishers should consider making their meta-data for publication timelines freely available. For example, in order to inform potential authors eLIFE, a non-profit organization that operates a publishing platform for scientists, provides immediate and downloadable access to the platform’s submission volume and publication timeline data in their author instructions [19]. Journal publishers might go a step further and join data sharing consortia that allow for standardization, aggregation, and dissemination of this data. Analysis of this data might highlight exemplar journals with efficient publication timelines, the best practices of which other journals could emulate.

## Conclusion

In this study, we used publicly available data to determine publication timelines in HPE journals that make this information available. Our data, while incomplete, suggest that HPE researchers may face longer timelines than their counterparts in the biomedical sciences. Perhaps more important than this finding was our ability to use Open Science and scientific replication to answer, albeit incompletely, an important question in HPE. As a next step, we call on all HPE journals to consider sharing their publication timeline data with transparency and completeness so that all stakeholders in the publication process can have access to accurate, open information.

